# Different effects of 3-week disuse on phenotype and gene expression in calf and thigh muscles

**DOI:** 10.64898/2025.12.04.691987

**Authors:** Mira A. Orlova, Nadia S. Kurochkina, Roman Yu. Zhedyaev, Nikita E. Vavilov, Viktor G. Zgoda, Alina A. Puchkova, Anna A. Borzykh, Tatiana F. Vepkhvadze, Egor M. Lednev, Pavel A. Makhnovskii, Gleb V. Galkin, Valery E. Novikov, Alexey V. Shpakov, Daniil V. Popov

## Abstract

Disuse, like several other pathological conditions, has specific effects on various skeletal muscles; the mechanisms underlying these responses remain unclear. We aimed to compare the disuse-induced changes in the phenotype and proteome of the calf and thigh muscles, and to assess the extent to which these proteomic changes are regulated at the mRNA and other levels. Twelve healthy young males participated in 3-week bed rest. Disuse resulted in a greater decrease in lean mass, aerobic performance, and changes in the proteome and transcriptome of the calf muscles/*m. soleus* than the thigh muscles/*m. vastus lateralis*. A greater decrease in calf muscle mass was associated with a decrease in the expression/deactivation of translation regulators, but not to the expression of the main sarcomeric proteins. At the same time, a significant decrease in aerobic performance of the ankle plantar flexors occurred without changing the expression of oxidative enzymes – a marker of mitochondrial density. That decrease was associated with dysregulation of mitochondrial biogenesis. Most large-scale changes in the transcriptome did not translate into changes in the proteome, indicating post-transcription protein buffering. However, changes in the RNA levels were revealed to play a dominant role in regulating specific proteins, whereas for others, this factor played little or no role. In conclusion, our findings partially explain why calf muscles with a strong postural function are more sensitive to short-term disuse. This provides a foundation for developing targeted approaches to counteract the negative effects of disuse on different muscles.

## INTRODUCTION

Disuse lasting more than a week causes a significant decrease in mass and functional capacity of skeletal muscles, including maximum voluntary contraction, aerobic performance, the ability to oxidize fats and carbohydrates, and insulin sensitivity (Mikines *et al*., 1991; Ried-Larsen *et al*., 2017; Marusic *et al*., 2021). A decrease in muscle mass is mainly observed in the postural trunk and leg muscles; moreover, the calf muscles, in particular *m. soleus*, show the greatest decrease (Hardy *et al*., 2022). This is mainly due to the fact that *m. soleus* has a strong postural function and is composed of more than 80% slow-twitch type I muscle fibers. It is much more sensitive to a reduction in contractile activity than other leg muscles, in terms of muscle mass, fiber size, maximum voluntary contraction, and tone (LeBlanc *et al*., 1992; Kozlovskaya *et al*., 2007; Fitts *et al*., 2010; Tomilovskaya *et al*., 2019; Casuso *et al*., 2024; Borzykh *et al*., 2025b). The loss of skeletal muscle mass and functional capacity during disuse have been studied for decades, but the mechanisms underlying the specific responses of different muscles to disuse remain unclear.

A key factor regulating the disuse-induced decrease in the mass of human “mixed” skeletal muscles (e.g., *m. vastus lateralis*) is a decrease in translation rate (Crossland *et al*., 2019; Brook *et al*., 2022; Deane *et al*., 2024). However, it is unclear whether this decrease varies between different muscles. Although literature has described the possibility of transcript-specific regulation of individual mRNA translation (Leppek *et al*., 2018), the disuse-induced decrease in protein synthesis has a generalized effect that appears to suppress the translation of most or all mRNAs. This should lead to a decrease in the absolute amount of most or all proteins in the muscle fibers (and a decrease in muscle mass) without changing proteins concentration. On the other hand, the rate of mRNA translation (and the number of synthesized protein molecules) is regulated by changes in mRNA concentration. Interestingly, after 6 days of disuse, the expression of three times more genes (~1,500 mRNAs) changed in human *m. soleus* than in *m. vastus lateralis* (~600 mRNAs) (Borzykh *et al*., 2025a; Borzykh *et al*., 2025b), and this difference persisted after 2 months of disuse (Chopard *et al*., 2009). It means that disuse may lead to appropriate changes of proteins concentration in *m. soleus* and *m. vastus lateralis* that partially explained the difference in decreased functional capacity of these muscles.

In human skeletal muscles, changes in the proteome profile induced by disuse occur more slowly than changes in the transcriptome. Indeed, after 3 days (*m. soleus*) and 6 days (*m. soleus* and *m. vastus lateralis*) of “dry” immersion, no changes were detected (Popov *et al*., 2023; Borzykh *et al*., 2025b), whereas after 14 days of limb immobilization, only 99 proteins in *m. vastus lateralis* changed their concentrations (Doering *et al*., 2022). This is because *i*) in these studies less than one-third of the skeletal muscle proteome were identified, *ii*) changes in gene expression (mRNA levels) are compensated at the post-transcriptional level, which is aimed at preventing proportional changes in the levels of specific proteins (post-transcription protein buffering) (Heller *et al*., 2026), or *iii*) most of highly abundant proteins have a half-life of more than one week, in particular cytoskeletal/sarcomeric proteins, extracellular matrix and mitochondrial proteins, and histones (Fornasiero *et al*., 2018; Rolfs *et al*., 2021). The latter suggests that the changes in the concentration of many mRNA occurred in the first days of disuse may lead to changes in the protein concentration and functional capacity of muscles at a later stage. These changes are likely to be more pronounced in *m. soleus* compared to *m. vastus lateralis*.

The aim of the study was: *i*) to compare the changes in the phenotype and proteomic profile of the ankle plantar flexor muscles/*m. soleus* and knee extensors/*m. vastus lateralis* after three weeks of head-down bed rest, and *ii*) to assess the extent to which these proteomic changes are regulated at the mRNA and other levels. For this purpose, the mass of the thigh and calf muscles, their maximum voluntary contraction, and aerobic performance were measured before and after disuse, and compared to the maximum mitochondrial respiration rate, proteomic (global quantitative mass spectrometry-based proteomic) and transcriptomic (RNA-sequencing) profiles of samples from *m. soleus* and *m. vastus lateralis* (Fig. 1).

**Fig. 1.**
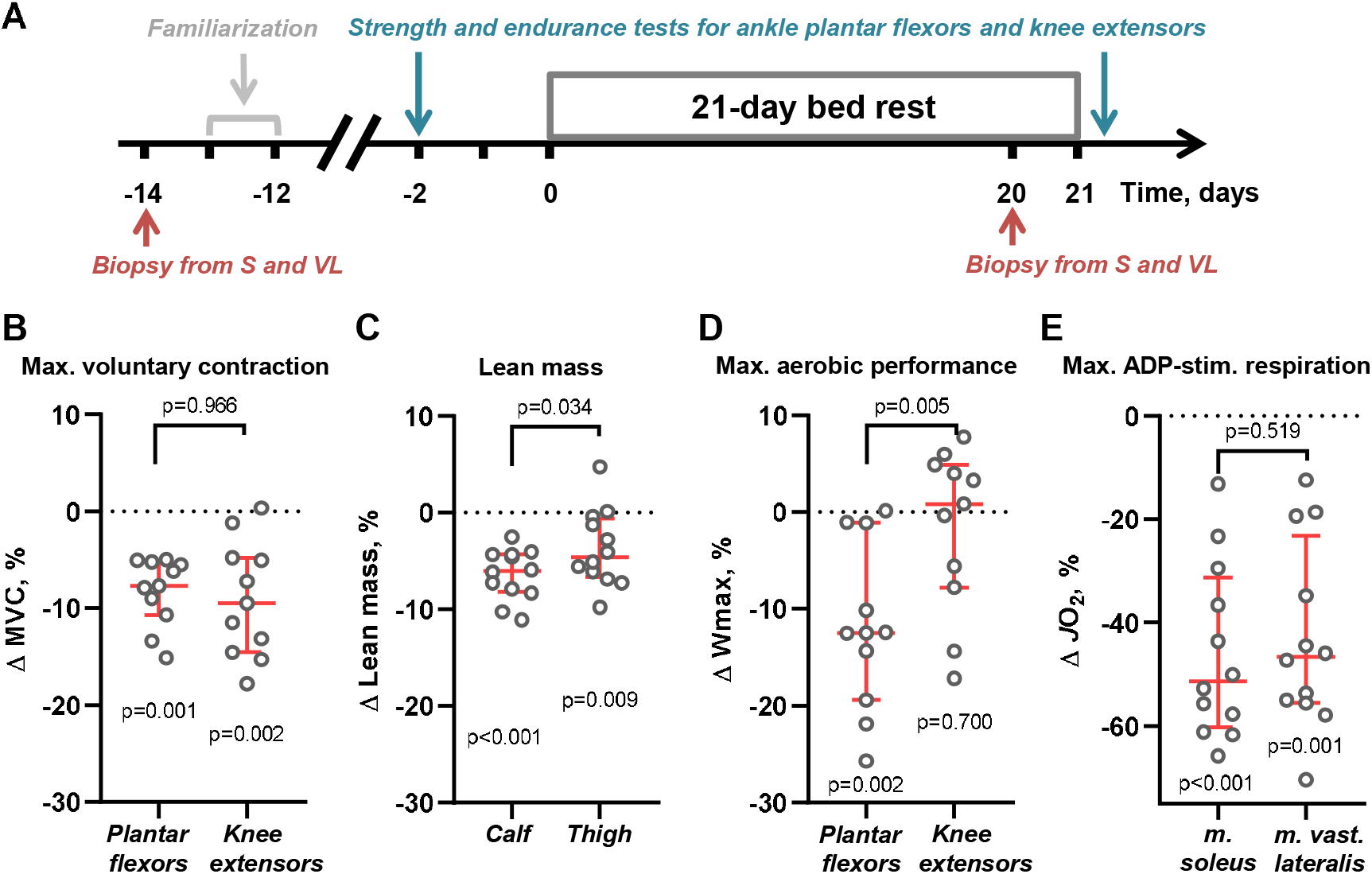
Three-week bed rest led to more pronounced changes in the phenotype of calf muscles compared to thigh muscles. **(A)** Study design; *m. soleus* (*S*) and *m. vastus lateralis* (*VL*) (n=12 subjects). **(B)** Changes in the maximum isometric voluntary contraction of the ankle plantar flexors and knee extensors (n=11 subjects). **(C)** Changes in calf and thigh lean mass (n=12 subjects). **(D)** Changes in the maximum power output of the ankle plantar flexors and knee extensors in the incremental ramp test (n=11 subjects). **(E)** Changes in maximum ADP-stimulated mitochondrial respiration (*JO*_*2*_) in permeabilized fibers of *m. soleus* and *m. vastus lateralis* (n=12 subjects).

## METHODS

### Ethical approval

The study was approved by the Biomedical Ethics Committee of the Institute for Biomedical Problems of the RAS (Protocols No. 599, October 6, 2021, and No. 621, August 8, 2022) in compliance with the Declaration of Helsinki. All volunteers provided written informed consent for participation and skeletal muscle biopsy.

### Study design

Design of the study was described previously (Puchkova *et al*., 2024). Briefly, 12 healthy men (aged 24–40 years; median height: 1.78 m interquartile range [1.73–1.82 m]; body mass: 78 kg [72–81 kg]; BMI: 23.4 kg/m^2^ [22.3–26.9 kg/m^2^]) participated in a 21-day head-down (-6 degree) bed rest. Light-off period was set at 23:00-07:00. Subjects were constantly monitored with video. The average time spent on daily hygiene on odd days (without a shower) was <7 min, and on even days (with a shower) <15 min; for day 21, it was <50 min due to muscle biopsy. To minimize the time spent in a vertical position, the subjects were transferred and took a shower in a horizontal position. Dietary intake was controlled during the study; the menu composition was identical for all subjects.

Following a familiarization session, muscle performance testing was conducted two days prior to and 8-10 h after the bed rest (after other tests, which were not included in the study) (Fig. 1A). The maximum isometric voluntary contraction (MVC) of the ankle plantar flexors and knee extensors in the right leg was examined. After a 10-minute rest period, aerobic performance of these muscle groups was measured (the maximum power [*W*_*max*_] in an incremental ramp test to exhaustion). Muscle biopsies were obtained using a Bergstrom needle under local anesthesia (2 ml of 2% lidocaine) from the middle part of *m. vastus lateralis* and *m. soleus* 14 days prior to and 20 days following the start of bed rest (Fig. 1A). Biopsies were performed at 10:00 am, three hours after a standardized light breakfast (5.2 g protein, 2.7 g fat, 55 g carbohydrates, 1253 kJ). Samples were processed to remove connective and adipose tissue; part of the fresh tissue was used to assess mitochondrial function; another part was immediately frozen in liquid nitrogen to study the transcriptome and proteome profiles; all frozen samples were stored at -80°C until further analysis.

### Lean mass

The lean mass of each leg and calf was assessed using dual-energy X-ray absorptiometry (scan mode “Whole Body”, densitometer Hologic, USA). To calculate the thigh lean mass, the calf lean mass was subtracted from the leg lean mass. Mean values for the left and right legs were then calculated.

### Muscle performance tests

All testing protocols were described elsewhere (Borzykh *et al*., 2025b). Briefly, to reduce the contribution of the biarticular gastrocnemius muscles to force production, isometric MVC of plantar flexors was assessed using a CalfRaise-PRO dynamometer (AntexLab, Russia) in a sitting position (knee joint angle was 70 degrees, position of the calf was vertical, and the ankle joint angle was 90 degrees; the distal part of the foot was placed on a 4 cm high stand) (Fig. A1). Then, the isometric MVC of the knee extensors was examined in a sitting position at a knee joint angle 110 degrees using a Pro System 3 dynamometer (Biodex, USA). In each test, each participant completed 5 attempts, separated by at least a 30-second rest period; the best one was taken into account.

Maximum aerobic power (*W*_*max*_) of the ankle plantar flexors was assessed using a CalfRaise-PRO ergometer in an incremental ramp test to exhaustion in a sitting position (as described above). The initial load for raising the calf was 30 N, the load increment was 25 N/min; the load during calf lowering was minimal. The subject performed rhythmic (0.5 cycle/s) flexions and extensions of the ankle joint (raising the calf with 55% of the maximum amplitude) until they could no longer maintain the full range of motion for five cycles in the given rhythm.

The *W*_*max*_ of the knee extensors was assessed with an incremental ramp test to exhaustion using a modified Ergoselect 900 cycle ergometer (Ergoline, Germany) in a reclining sitting position (hip joint 135 degrees) (Fig. A1). Each subject performed rhythmic (1 cycle/s) knee extensions and flexions (from 80 to 160 degrees). The initial load was 0 W, the load increment – 1.1 W/min, and the load during knee flexion was minimal. The test was stopped when the subject was unable to maintain the full amplitude for 5 cycles in the given rhythm.

### Mitochondrial respiration in permeabilized muscle fibers

As described previously (Borzykh *et al*., 2025b), a piece of fresh tissue (~10 mg) was placed in ice-cold BIOPS relaxation buffer (10 mM Ca-EGTA buffer, 0.1 μM free calcium, 20 mM imidazole, 20 mM taurine, 50 mM K-MES, 0.5 mM DTT, 6.56 mM MgCl_2_, 5.77 mM ATP, 15 mM phosphocreatinine, *pH* 7.1) to remove connective and adipose tissue. Then a small piece (~2 mg) was removed and the muscle fibers were separated using a pair of needles to create a fiber bundle. The fibers were permeabilized in BIOPS buffer containing saponin (50 μg/ml, 30 min) on ice with slow stirring, then washed twice for 10 minutes in MiRO5 buffer (0.5 mM EGTA, 3 mM MgCl_2_·6H_2_O, 60 mM lactobionic acid, 20 mM taurine, 10 mM KH_2_PO_4_, 20 mM HEPES, 110 mM D-sucrose, 1 g/l fatty acid-free bovine serum albumin). Mitochondrial respiration rate was measured simultaneously on two separate channels using an Oxygraph polarograph (Hansatech, UK) under hyperoxic conditions (*Co*_*2*_ >200 μM). Substrates were added sequentially: malate (2 mM; final concentration) + pyruvate (5 mM) + glutamate (10 mM), ADP (5 mM) + MgCl_2_ (3 mM), succinate (10 mM) (to examine maximal ADP-stimulated respiration, state 3), and cytochrome C (10 μM) (to check mitochondrial membrane integrity). The respiration rate without the addition of substrates was subtracted from the respiration rate after the addition of substrates, then the respiration rate was normalized to the mass of permeabilized muscle fibers.

### Quantitative mass spectrometry-based proteomics and data processing

As described previously (Kurochkina *et al*., 2024), a piece of frozen tissue (~10 mg) was homogenized in 140 μl of lysis buffer (4% sodium dodecyl sulfate, 0.1 M Tris and 0.1 M dithiothreitol, *pH* 7.6). The lysate was heated (95ºC, 5 min), transferred to an AFA microtube, sonicated (average power 20 W, 30 s × 4) using a ME220 sonicator (Covaris, USA), and centrifuged (5 min, 30,000 g). The protein concentration in the supernatant was measured by a fluorimeter Qubit 4.

The proteins (85 μg) were diluted in lysis buffer (final volume 24 μl), transferred to an S-Trap column (ProtiFi, USA), alkylated (20 mM iodoacetamide, 15 min), and hydrolyzed into the S-Trap column using trypsin (2 h at 47°C, 1:10, Promega, USA) in 40 μl 50 mM triethylammonium bicarbonate according to the manufacturer’s recommendations. After elution, 20 μg of peptides (fluorimeter Qubit 4) were dried, resuspended in 100 mM triethylammonium bicarbonate, and labelled with TMT 16-plex isobaric labels (Thermo Scientific, USA) for 1.5 h according to the manufacturer’s recommendations. The reaction was stopped (0.3% hydroxylamine w/v, 15 min), and samples were pooled.

The mixture of labelled peptides was separated using high *pH* reverse-phase LC fractionation (HPLC 1200, Agilent, USA). The peptides were concentrated on an analytical column (XBridge C18, particle size 5 μm, 4.6 × 250 mm, Waters, Ireland) in isocratic mode at a flow of 750 μl/min for 3 min in mobile phase A (15 mM ammonium acetate, *pH* 9.0). Twenty-four fractions were collected from 3 min to 50 min (collection time 2 min, volume 1500 μL) using a gradient elution mode with mobile phase B (15 mM ammonium acetate, *pH* 9.0, 80% acetonitrile, *pH* 9.0). Each fraction was concentrated, and then fractions (1 and 13, etc.) were combined to obtain 12 resulting fractions.

Each fraction was analyzed two times using an HPLC Ultimate 3000 RSLC nanosystem (Thermo Scientific, USA) and a Q Exactive HF-X Hybrid Quadrupole-Orbitrap mass spectrometer (Thermo Scientific, USA) by the nanoelectrospray ion source in the positive mode of ionization (Thermo Scientific) as previously described. The gradient (60 min) was formed by the mobile phase A (0.1% formic acid) and B (80% acetonitrile, 0.1% formic acid) at a 0.33 μL/min flow. The ionizing voltage was 2.1 kV. MS spectra were acquired at a resolution of 60,000 in the 440–1800 m/z range; fragment ions were mass scanned at a resolution of 60,000 at the range from m/z 120 to the upper m/z value as assigned by the mass to charge state of the precursor ion. All tandem MS scans were performed on ions with a charge state from z=2+ to z=4+. Synchronous precursor selection facilitated the simultaneous isolation of up to 20 MS2 fragment ions. The maximum ion accumulation times were set to 50 ms for precursor ions and 25 ms for fragment ions. AGC targets were set to 10^6^ and 10^5^ for precursor ions and fragment ions, respectively.

Peptide and protein identification and search were conducted using the MaxQuant platform (version 2.1.4.0; Max Planck Institute for Biochemistry, RRID:SCR_014485) using default settings (FDR for peptides 1%, N-terminal acetylation and oxidation of methionine as variable modifications, and carbamidomethylation of cysteine, as fixed modification) and the *Isobaric much between runs* and *PSM-level weighted ratio normalization* functions (Yu *et al*., 2020). Further analysis was performed using a Perseus platform (1.6.5; Max Planck Institute for Biochemistry, RRID:SCR_015753). After filtration (potential contaminants, reverse peptides, peptides identified only by site), proteins identified by more than one peptide (unique+razor) and presented in 100% of the samples were selected for further analysis. Reporter ion intensities were log-transformed and normalized to the median. To compare response to disuse in different muscles the 2-way ANOVA (interaction of factors Muscle × Disuse) was used (Perseus platform); *p*-values were adjusted (FDR) using the *p*.*adjust* package (R environment). The effect of disuse on each muscle was assessed using the two-sample Welch T-test with permutation-based FDR (*q*- or *p*_*adj*_-value; Perseus platform). Differentially expressed proteins were defined using *p*_*adj*_ <0.05.

To investigate the influence of muscle composition on the magnitude of change in protein expression after disuse the expression (unit proportional to molar concentration) of myosin heavy chain isoform MYH7 (as well MYH1 and MYH2) was calculated in each sample (*i*) as:

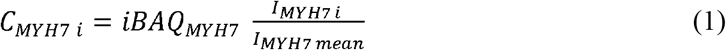

where, *i BAQ*_*MYH7*_ – Intensity-Based Absolute Quantification, the mean expression of the MYH7 isoform in all samples, *I* ^*MYH7i*^ – the reporter ion intensity of the MYH7 isoform in sample *i, I*_*MYH7 mean*_ – the median reporter ion intensity of the MYH7 isoform in all samples.

Then, Spearman correlations between the proportion in expression of the MYH7 isoform and disuse-induced changes in protein expression were exanimated separately in each muscle using *p*<0.05.

### Western blot analysis

As described previously (Borzykh *et al*., 2025b), the lysate samples were mixed with Laemmli buffer and loaded onto 10% polyacrylamide gels (20 μg protein/lane). Electrophoresis (20 mA per gel) was performed using a Mini-PROTEAN Tetra Cell system (Bio-Rad, USA). The proteins were transferred to nitrocellulose membranes using a Trans-Blot Turbo system (Bio-Rad) in Towbin buffer for 30 min at 25 V. The membranes were stained with Ponceau S to verify consistent loading of proteins, followed by washing and incubation for 1 h in 5% non-fat dry milk (Sigma-Aldrich, USA) in TBST (150 mM NaCl, 0.1% Tween 20, 20 mM Tris-HCl, *pH* 7.6). Next, the membranes were incubated at 4°C overnight with antibodies: anti-phospho-4EBP1 (Eukaryotic translation initiation factor 4E-binding protein 1) Thr37/46 (1:1000; 236B4; #2855, RRID:AB_560835) or anti-phospho-EF2 (Elongation factor 2) Thr57 (1:1000; #2331, RRID:AB_10015204) (all from Cell Signaling Technology, USA) or anti-phospo-KS6A1 or p90 (Ribosomal protein S6 kinase alpha-1) S380 (1:1000, ab32203, RRID:AB_777767) or anti-G3P or GAPDH (Glyceraldehyde-3-phosphate dehydrogenase; 1:2500, ab9485, RRID:AB_307275) or anti-TBA4A (Tubulin alpha-4A chain; 1:7500, ab176560, RRID:AB_2860019) (all from Abcam, UK) in 5% non-fat dry milk in TBST. Membranes were then incubated for 1 h at room temperature with an anti-rabbit IgG secondary antibody (1:10000, #7074, Cell Signaling Technology, RRID:AB_2099233) in 5% non-fat dry milk in TBST. After each step, the membranes were washed three times for 5 min with TBST. Finally, the membranes were incubated with SuperSignal West Femto Maximum Sensitivity Substrate (Thermo Fisher Scientific, USA) and luminescent signals were captured using a ChemiDoc Imaging System (Bio-Rad). Images were analyzed using the Image Lab Software (v6.0.1, Bio-Rad, RRID:SCR_014210); signals from phosphoproteins were normalized to the mean signal from G3P and TBA4A.

### RNA sequencing and data processing

A frozen tissue sample (~15 mg) was homogenized at 4°C in 500 μl of a lysis buffer (LRU-100-50, Biolabmix, Russia) using a plastic pestle and a drill (200 rpm, ~ 30 s). Total RNA (without small RNA fraction) was extracted with phenol-chloroform method and selective sorption on a silicon membrane. RNA concentration was measured with a Qubit 4 fluorometer (ThermoScientific, USA), and RNA integrity was measured using capillary gel electrophoresis (TapeStation, Agilent, USA). The median RNA integrity number (RIN) across all samples was 8.2 (7.9-8.3).

As described previously (Kurochkina *et al*., 2024), strand-specific RNA libraries were prepared using 500 ng of total RNA, by the NEBNext Ultra II RNA kit (New England Biolabs, USA) in accordance with the manufacturer’s instructions. The efficient concentration of the libraries was assessed by qPCR using the 5X qPCRmix-HS SYBR kit (Evrogen, Russia). Single-end sequencing was performed on a NextSeq 550 analyzer (Illumina, USA) with a read length of 75 bp and an average of ~60 million reads per sample. Raw data has been deposited to NCBI GEO: GSE310858.

Sequencing quality was assessed using the *FastQC* tool (v.0.11.5, RRID:SCR_014583), and low-quality reads were removed using *Trimmomatic* (v.0.36, RRID:SCR_011848). High-quality reads were mapped to the human reference genome GRCh38.p13 primary assembly. The number of unique reads mapped to exons of each gene was determined using the *Rsubread* package (R environment) and Ensembl annotation (GRCh38.101). Protein-coding genes (including polymorphic pseudogenes and translated pseudogenes) with TPM >1 (Transcripts Per Million, kallisto tool v0.46.2) were used to analyze differential gene expression by the DESeq2 method using the paired Wald test (with factor Subject) and Benjamini-Hochberg correction. The influence of different muscles on the response to disuse was assessed using a model with interaction of factors (Muscle × Disuse) for paired comparisons (Wald test). Differentially expressed genes were defined as protein-coding genes with *p*_*adj*_ <0.01, and magnitude of change ≥25%.

### Statistical analysis

Data are presented as the median and interquartile range. A nonparametric Wilcoxon test was used to evaluate the significance of changes following disuse or between different muscle groups, with a significance level set at 0.05. For transcriptomic and proteomic data, a functional enrichment analysis for biological processes, cellular components and pathways (UNIPROT KW BP/CC, GENE ONTOLOGY BP/CC DIRECT and KEGG PATHWAY databases) was performed against all expressed protein-coding genes (TPM >1, ~11,000 mRNA) or detected proteins (~1,900 proteins) using the DAVID tool v2023q4 (RRID:SCR_001881) (Fisher’s exact test with Benjamini correction, *p*_*adj*_ <0.05). Forward stepwise multiple regression was used to examine the effect of protein half-life and changes in mRNA levels on disuse-induced changes in protein expression; mouse skeletal muscle protein half-life data were taken from (Rolfs *et al*., 2021).

## RESULTS

### The functional capacity of calf and thigh muscles

Three weeks of bed rest resulted in a comparable reduction in maximal voluntary contraction (MVC) between the ankle plantar flexors (median 7.7% and interquartile range (5.2 - 10.7)%, *p*=0.001) and knee extensors (9.4(4.8 - 14.6)%, *p*=0.002) (Fig. 1B, Fig. A2). However, the decrease in lean mass was greater (*p*=0.034) in the calf, than in the thigh (6.0(4.0 - 8.0)%, *p*<0.001 and 4.5(0.2 - 6.7)%, *p*=0.009, respectively) (Fig. 1C, Fig. A2). In addition, a decrease in maximal aerobic power was only observed in the plantar flexors (12.5(1.1 - 19.4)%, *p*=0.002); moreover, this response was greater than in the knee extensors (*p*=0.005) (Fig. 1D, Fig. A2). Meanwhile, the decrease in maximal ADP-stimulated mitochondrial respiration in permeabilized fibers was comparable in *m. soleus* and *m. vastus lateralis* (51(31 - 60)% and 47(23 - 55)%, respectively, both *p*≤0.001; Fig. 1E, Fig. A2).

The median and interquartile range. Dotted line – the level prior to disuse. Lower symbols – effect of disuse; upper symbols – differences in response to disuse between muscles (Wilcoxon test).

### Proteomic profile in *m. soleus* and *m. vastus lateralis*

Across all samples, 1,844 proteins were identified, predominantly, highly abundant mitochondrial and cytoskeletal/sarcomeric proteins (Fig. A3). Principal component analysis revealed that the differences in proteome between the muscles were more pronounced than the effects of bed rest (Fig. A4A). However, after bed rest, there were noticeable differences between the muscles: 170 proteins changed expression in *m. soleus*, whereas only 32 – in *m. vastus lateralis*. These responses significantly overlapped, with 5 proteins responding only in *m. vastus lateralis* (Fig. 2A, B, Supporting Information Table S1). Proteins that responded in both muscles had a comparable magnitude of expression change. However, for proteins that only responded in *m. soleus*, the magnitude was twice as large as in *m. vastus lateralis* (Fig. A4B). Despite this, no interaction between factors Muscle × Disuse was found (*p*_*adj*_ >0.431), apparently due to the large intragroup variation (Fig. 2B).

**Fig. 2.**
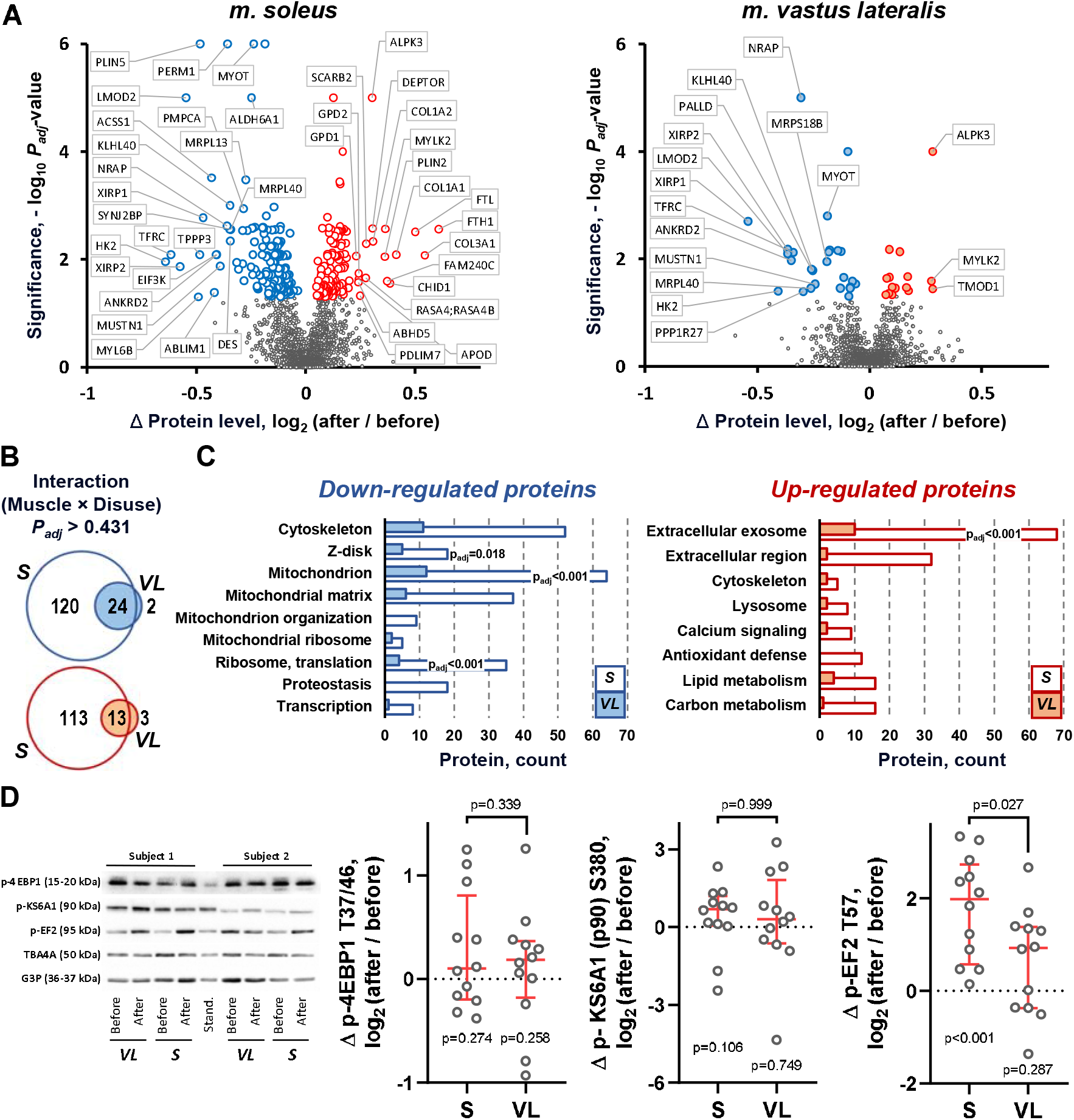
Three-week bed rest led to greater changes in the proteomic profile in *m. soleus* (*S*) compared to *m. vastus lateralis* (*VL*) (n=11 subjects). **(a)** The distribution of the expression changes: blue/red points show down- and up-regulated proteins (*p*_*adj*_ <0.05), respectively; proteins with a greater response are indicated (see also Supporting Information Table S1). **(b)** The proteomic response to bed rest, in terms of the number of differentially expressed proteins (*p*_*adj*_ <0.05), significantly overlapped between *m. vastus lateralis* and *m. soleus* for both down- and up-regulated proteins. **(c)** Functional annotation of differentially expressed proteins. Enriched categories are indicated (see also Supporting Information Table S1). ***(d)*** Changes in the phosphorylation level of canonical sites in translation regulators. The median and interquartile range. Dotted line – the level prior to disuse. Lower symbol – effect of disuse; upper symbol – difference in response to disuse between muscles (Wilcoxon test). *Stand*. –a standard sample, a mixture of six samples of *m. soleus* lysates.

Expression of the major skeletal muscle myosin heavy chain isoforms (MYH1/2/7) were unchanged after disuse in *m. soleus* (*p*_*adj*_ = 0.646, 0.496, and 0.262, respectively) and *m. vastus lateralis* (*p*_*adj*_ = 0.810, 0.356, and 0.646, respectively) (Supporting Information Table S1), as well as their proportion (Fig. A7). A significant correlation between the proportion in expression of the MYH7 isoform and disuse-induced changes in protein expression in both muscles was found for 18 proteins. Interestingly, only 4 of these proteins showed significant changes in expression after bed rest in both *m. soleus* and *m. vastus lateralis* (Unconventional myosin MYO18A *r*_*s*_ = - 0.62, *p*_*adj*_ = 0.043 and *r*_*s*_ = -0.67, *p*_*adj*_ = 0.023; PDZ and LIM domain protein PDLIM7 *r*_*s*_ = 0.66, *p*_*adj*_ = 0.023 and *r*_*s*_ = 0.65, *p*_*adj*_ = 0.032; Phosphoglucomutase PGM1 *r*_*s*_ = 0.76, *p*_*adj*_ = 0.007 and *r*_*s*_ = 0.76, *p*_*adj*_ = 0.007; Triadin TRDN *r*_*s*_ = 0.64, *p*_*adj*_ = 0.035 and *r*_*s*_ = 0.61, *p*_*adj*_ = 0.047). This result suggests that intragroup muscle composition variation has little influence on the bed rest-induced changes in protein expression. This is indirectly related to the results of a meta-analysis showing that disuse causes a comparable decrease in the cross-sectional area of type I and II fibers in a muscle (Vikne *et al*., 2020).

Additionally, only a few significant correlations between changes in protein expression in both muscles and changes in phenotypic outcomes were found. This is likely because maximal voluntary contraction and maximal aerobic performance are partially dependent on neuromuscular control and were measured for a muscle group (knee extensors or ankle flexors), whereas lean mass was measured across all thigh or calf muscles.

In both muscles, bed rest decreased the expression of proteins included in the categories (UniProt, GO, and KEGG): ‘*cytoskeleton’*, ‘*Z-disk’*, ‘*mitochondrion’*, ‘*mitochondrial matrix’*, ‘*mitochondrial ribosome’*, ‘*ribosome, translation’*, and ‘*transcription’*; while up-regulated proteins were related to ‘*extracellular exosomes’*, ‘*extracellular region’*, ‘*cytoskeleton’*, ‘*lysosome’*, ‘*calcium signaling’*, ‘*lipid metabolism’*, and ‘*carbon metabolism’* (Fig. 2C). The response of *m. soleus* was substantially more pronounced in terms of the protein number and included specific categories: ‘*proteostasis’* and ‘*mitochondrial organization’* for down-regulated proteins, and ‘*antioxidant defense’* for up-regulated proteins. Moreover, only *m. soleus* demonstrated significant enrichment for certain categories: ‘*Z-disk’*, ‘*mitochondrion’*, ‘*ribosome, translation’*, and ‘*extracellular exosome’* (Fig. 2C).

### Translation regulation in *m. soleus* and *m. vastus lateralis*

Using western blotting, we found no changes in the phosphorylation levels of the canonical sites of Eukaryotic translation initiation factor 4E-binding protein 1 (4EBP1) ^T37/46^ and Ribosomal protein S6 kinase alpha-1 (KS6A1 or p90) ^S380^ in both muscles (Fig. 2D). However, in *m. soleus*, a substantial increase in the phosphorylation of another regulator, Elongation factor 2 (EF2) ^T57^ – a marker of its deactivation, was observed (p<0.001); moreover, this response was greater than in *m. vastus lateralis* (p=0.027 to 0.999) (Fig. 2D).

### Transcriptomic profile in *m. soleus* and *m. vastus lateralis*

After removing low-expressed genes, >11,000 mRNAs were identified (Supporting Information Table S2). Principal component analysis revealed that the transcriptomic profile of *m. soleus* was markedly different from that of *m. vastus lateralis* and the disuse-induced changes in the transcriptomic profile of *m. soleus* were substantially greater than those in *m. vastus lateralis* (Fig. A4C). Indeed, 2,018 genes were differentially expressed in *m. soleus*, whereas only 592 – in *m. vastus lateralis*, and most of these (70-80%) overlapped with those found in *m. soleus*. Furthermore, an interaction Muscle × Disuse was found for 920 mRNAs, confirming a stronger response of *m. soleus* to disuse (Fig. 3A; Supporting Information Table S2). Genes that were over-suppressed in *m. soleus* after disuse (relative to *m. vastus lateralis*) enriched in categories associated with mitochondrial biogenesis (>400 out of ~1000 mRNAs encoding mitochondrial proteins, including most enzymes of the Krebs cycle and oxidative phosphorylation, as well as mitochondrial assembly factors, matrix and membrane proteins). These over-suppressed genes were also associated with regulators of translation, lipid, and amino acid metabolism. Genes that were over-activated in *m. soleus* were related to membrane, extracellular, and cytoskeletal proteins, as well as the immune and inflammatory responses (Fig. 3B-C, Supporting Information Table S2). This finding is consistent with the results of the functional enrichment analysis of genes that respond to disuse in both muscles and in *m. soleus* only (Fig. A5).

**Fig. 3.**
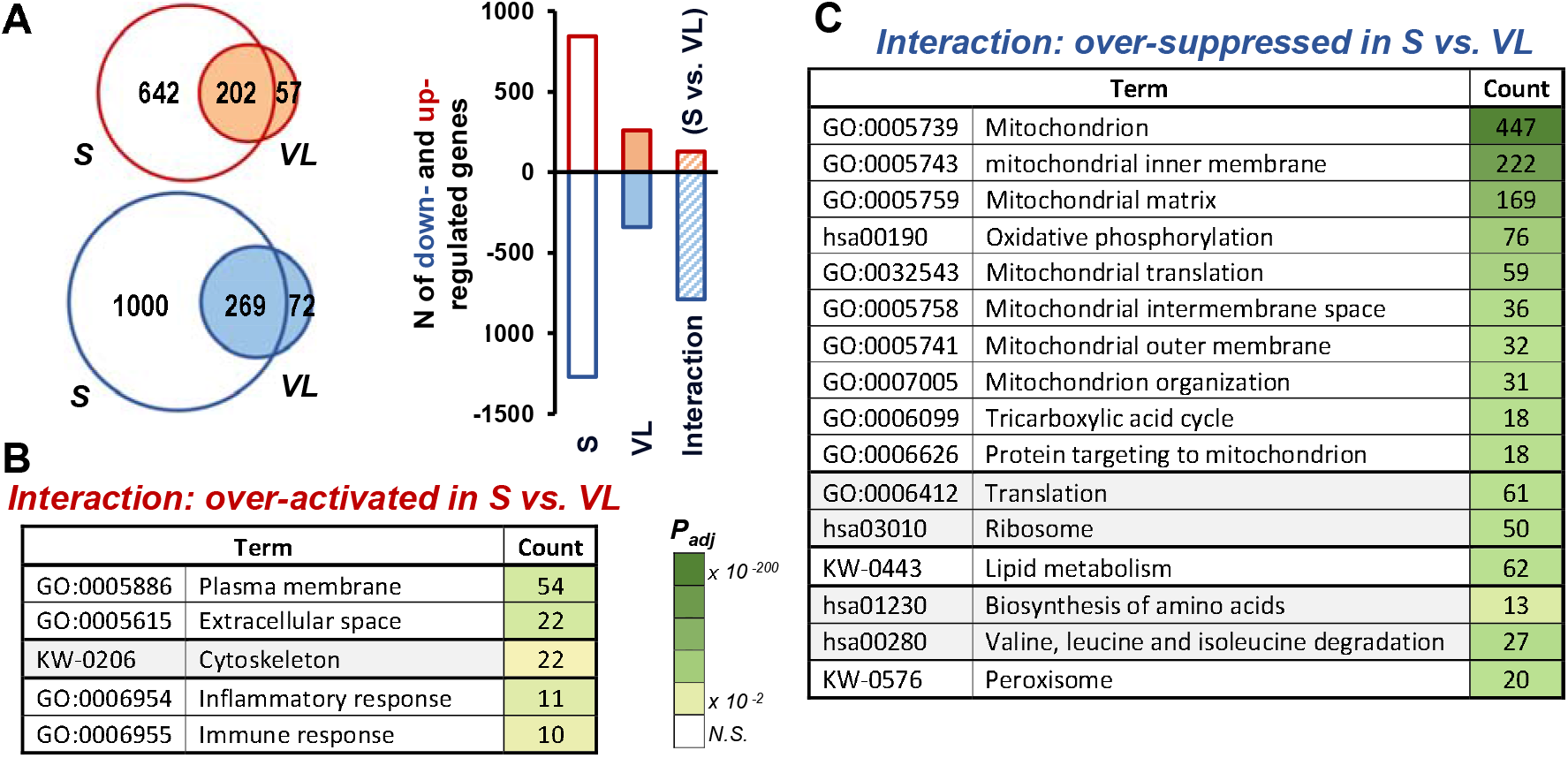
Three-week bed rest led to substantially greater changes in the transcriptomic profile of *m. soleus* (*S*) compared to *m. vastus lateralis* (*VL*) in terms of both the number of genes and the magnitude of the response (n=12 subjects). **(A)** The number of differentially expressed genes (*p*_*adj*_ <0.01, magnitude of change ≥25%) in *m. vastus lateralis* was substantially lower than in *m. soleus*. These down- and up-regulated genes were significantly overlapping. An interaction Muscle × Disuse was observed for 920 mRNAs, confirming a stronger response of *m. soleus* to disuse. **(B** and **C**) Functional enrichment analysis of differentially expressed genes that were over-activated or over-suppressed in S after disuse *vs*. VL. Heatmap shows the significance levels, the number – mRNA count (see also Supporting Information Table S2 and Fig. A5).

### Comparison of proteomic and transcriptomic responses

To determine how disuse-induced changes in protein expression are regulated at the mRNA levels, we compared the proteomic and transcriptomic responses and identified distinct patterns of regulation (Fig. 4A, Supporting Information Table S3). Multiple regression revealed that regulation at the mRNA level, as well as protein stability (data from (Rolfs *et al*., 2021)), exert significantly different effects on protein regulation across different regulatory patterns (Fig. 4B). Functional enrichment analysis revealed that in *m. soleus*, proteins with *‘mRNA Down - Protein Down’* regulation pattern were enriched in categories related to various mitochondrial proteins, particularly matrix proteins regulating mitochondrial biogenesis and proteostasis, but were not related to oxidative phosphorylation enzymes (Fig. 4C, Supporting Information Table S3). Meanwhile, proteins with the regulation pattern ‘*mRNA NS - Protein Down’* enriched in categories related to cytoplasmic proteins, including ribosomal and cytoskeletal (Z-disk) proteins, as well as regulators of focal adhesion, myogenesis, and differentiation. On the other hand, up-regulated proteins, showing no correlation with changes in their mRNAs (patterns ‘*mRNA NS - Protein Up’* and *‘mRNA Down - Protein Up’*), enriched in other categories, such as *‘extracellular exosome’* and *‘lipid metabolism’*. No significant enrichment was observed for *m. vastus lateralis*, due to the smaller number of differentially expressed proteins.

**Fig. 4.**
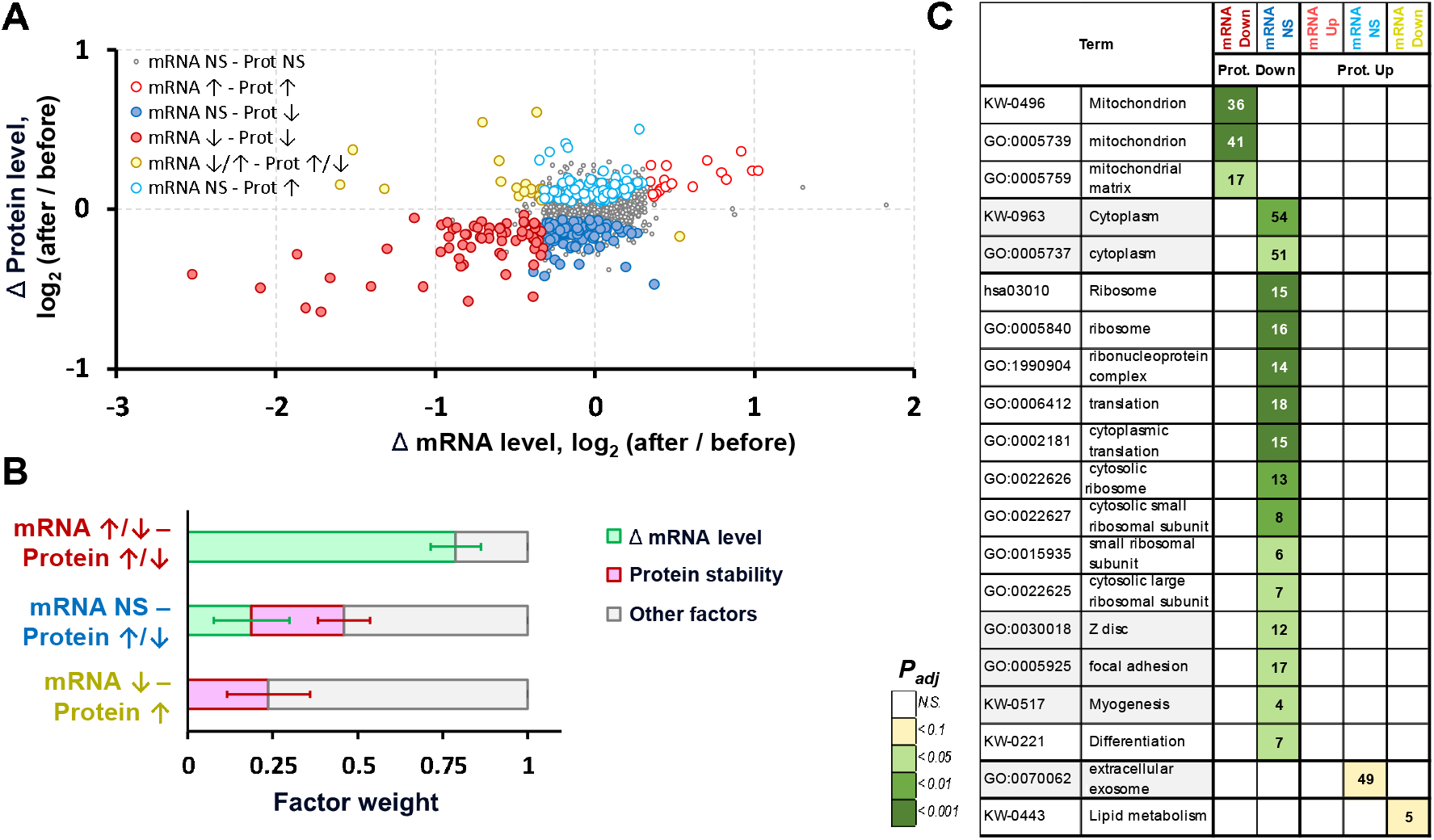
Patterns of protein regulation depend on their function. **(A)** Bed rest-induced changes in protein expression and the levels of corresponding mRNAs in *m. soleus* (each point depicts the median, different colors indicate different regulation patterns; *NS* – non significant). **(B)** The effect of changes in the mRNA levels and protein half-life on disuse-induced changes in protein expression for different regulatory patterns. Weight of multiple regression coefficient beta and standard error of beta are shown. **(C)** Functional enrichment analysis of proteins with different regulation patterns in *m. soleus*. The heatmap shows the significance levels, the number – mRNA number in enriched categories (see also Supporting Information Table S3). No enrichment for *m. vastus lateralis* was found.

Interestingly, in *m. soleus*, the transcriptomic analysis revealed substantial amounts of down-regulated mRNAs encoding most of glycolytic, Krebs cycle, and oxidative phosphorylation enzymes, as well as some regulators of lipid metabolism and muscle contraction (Fig. A5B). Although most of these proteins were identified by proteomic analysis, no changes in their expression were found (Fig. A6).

## DISCUSSION

### Three-week bed rest induced more pronounced changes in the phenotype of calf muscles compared to thigh muscles

Our study was the first to comprehensively compare the effects of short-term disuse (3 weeks) on the phenotype, proteome, and transcriptome of calf and thigh muscles. As expected, disuse led to a more pronounced decrease in lean mass in the calf (Marusic *et al*., 2021). Furthermore, a more pronounced reduction in aerobic performance was found in the ankle plantar flexor muscles compared to the thigh muscles (Fig. 1, Fig. A2). This supports the concept that muscles involved in maintaining posture (gravitationally dependent muscles) are more sensitive to reduced contractile activity (LeBlanc *et al*., 1992; Kozlovskaya *et al*., 2007; Fitts *et al*., 2010; Tomilovskaya *et al*., 2019; Borzykh *et al*., 2025b).

Despite the lack of a reduction in the aerobic performance of the knee extensors, the maximum ADP-stimulated mitochondrial respiration in permeabilized fibers of *m. vastus lateralis* decreased, and this decrease was comparable to that of *m. soleus*. Disuse did not cause changes in the expression of several dozen detected mitochondrial respiratory enzymes in both muscles (Fig. A6, Supporting Information Table S1). Together, these findings indicate a decrease in the intrinsic maximum ADP-stimulated mitochondrial respiration, apparently due to a dysregulation of mitochondrial respiration, which is consistent with the results of experiments with disuse of shorter duration (Miotto *et al*., 2019; Popov *et al*., 2023; Borzykh *et al*., 2025b).

### Potential mechanisms for a more significant change in the phenotype of *m. soleus*/ankle plantar flexors compared to *m. vastus lateralis*/knee extensors

Mass spectrometry-based proteomics enables the detection of predominantly highly abundant skeletal muscle proteins, such as sarcomeric, mitochondrial and ribosomal proteins, and chaperons; the coverage of low-abundant regulatory proteins is limited (Fig. A3). Our data, together with the results of other studies (3-, 6-, and 14-day disuse (Doering *et al*., 2022; Popov *et al*., 2023; Borzykh *et al*., 2025b) show that, in contrast to the transcriptome, changes in the expression of highly abundant proteins occur only after 2-3 weeks of disuse. The changes in the proteomic profile were significantly more extensive in “slow” *m. soleus*, in terms of the number of differentially expressed proteins, than in “mixed” *m. vastus lateralis*. This may explain, at least in part, the large difference in phenotype changes between calf and thigh muscles.

### Cytoskeletal and extracellular matrix proteins and regulators of protein synthesis

Despite the decrease in lean (muscle) mass of the calf and thigh, no decrease in the concentration of the main sarcomeric proteins was detected in both muscles (isoforms of myosin heavy chains encoded by *MYH1/2/7*, actin, tropomyosin, titin, nebulin, etc.). These proteins have the highest intensity of mass spectrometric peaks (10 out of top 15 proteins) (Fig. A7, Supporting Information Table S1), i.e., they constitute the largest mass portion of muscle proteins. Sarcomeric proteins with decreased expression in both muscles (encoded by *OBSCN, MYOT, PALLD, LMOD2*) or only in *m. soleus* (an unspecific for human skeletal muscle isoform of the myosin heavy chain, encoded by *MYH14*, as well as other sarcomeric proteins: *DES, MYOM1, MYL6B*/*12A*/*12B*/*18A, FLNB*/*C, ABLIM1, NEXN, LMOD3*) had a concentration two orders of magnitude lower than the main sarcomeric proteins (Fig. A7, Supporting Information Table S1). Therefore, the decrease in their expression does not explain the decrease in muscle mass and/or the greater decrease in mass in the calf muscles. However, the decrease in the expression of some Z-disk proteins found in both muscles (Fig. A7, Supporting Information Table S1) may be associated with a decrease in sarcomere transverse stiffness (Ogneva *et al*., 2009) that partially explains the decrease in MVC in both muscle groups.

A decrease in lean mass of the calf and thigh, occurring without a change in the concentration of the main sarcomeric proteins, indicates a proportional decrease in protein absolute content. This is apparently associated with a decrease in the rate of protein synthesis on ribosomes – the main mechanism for the decrease in the mass of “mixed” human muscle during short-term disuse (Crossland *et al*., 2019; Brook *et al*., 2022; Deane *et al*., 2024). The greater decrease in lean mass in the calf muscles, compared to the thigh muscles, may be explained by a more pronounced decrease in the rate of protein synthesis. This is indirectly confirmed by an increase in phosphorylation of the key elongation regulator EF2 ^T57^ only in *m. soleus* (Fig. 2D, Fig. A8). Additionally, in *m. soleus* there was a decrease in the expression of several subunits of translation initiation factors (encoded by *EIF3A*/*3J*/*3K*/*4B* genes) and four dozen ribosomal proteins (including mitochondrial ribosomal proteins) – a marker of a decrease in ribosome content (Fig. 2D, Supporting Information Table S1).

Interestingly, an increase in the expression of some cytoskeletal and extracellular matrix proteins was observed only in *m. soleus*, such as collagens (encoded by *COL1A1*/*1A2*/*3A1*/*15A1*), laminins (*LAMA2*/*B1*), sarcoglycan *SGCA*, and dystrophin (Fig. A7, Supporting Information Table S1). Fibrillar collagens types I and III that make up half of the extracellular matrix proteins (McKee *et al*., 2019) demonstrated some of the largest amplitudes of expression changes (increases of 1.24-to 1.53-fold). These changes in *m. soleus* are reminiscent of fibrotic changes in skeletal muscle associated with various pathological conditions (Mann *et al*., 2011; McKee *et al*., 2019; Agarwal *et al*., 2025). In addition, an increase in the expression of some proteins of the dystroglycan complex has already been noted after 84 days of disuse, both in *m. soleus* and in *m. vastus lateralis* (Chopard *et al*., 2005). Our results show that in *m. soleus* these changes occur significantly earlier, indicating greater sensitivity of this muscle to disuse. The increase in the concentration of key extracellular matrix proteins discussed above should result in an increase in the proportion of connective tissue in *m. soleus* and musculotendinous stiffness of the ankle plantar flexor muscles. This is confirmed by changes observed during longer-term conditions such as 60- and 90-day disuse (Haus *et al*., 2007; Thot *et al*., 2021) and space flight lasting 90 to 180 days (Lambertz *et al*., 2001), as well as age-related changes (Pavan *et al*., 2020).

### Proteostasis regulators

Disuse has been shown to reduce expression or alter phosphorylation of individual heat shock proteins (encoded by *Hspb1* [small heat shock protein], *Hspa1a*/*b* and *HSPA9* [HSP70 family]) and the transcription factor HSF1 in skeletal muscle of rodents (Lawler *et al*., 2006; Senf, 2013) and humans (Gram *et al*., 2014). In contrast to *m. vastus lateralis*, there was a large-scale decrease in the expression of heat shock proteins and their regulators in *m. soleus*: 23% or 18 of 79 detected proteins (Fig. 2C, Supporting Information Table S1). These proteins were localized both in the cytoplasm, e.g., those associated with families of the small heat shock protein (encoded by *CRYAB*), HSP40 (*DNAJA4*/*B4*), HSP70 (*HSPH1, BAG2*/*3*), and HSP90 (*HSP90AA1*/*AB1, AHSA1*), as well as *PFDN2*/4 and in mitochondria, e.g., those associated with the HSP10 (*HSPE1*), HSP60 (*HSPD1*), HSP70 (*HSPA9*), HSP90 (*HSP90AA1*/*AB1, TRAP1*), and *PFDN2*/*4* families. Studies with overexpression of *Hspb1, Hspa1a*/*b*, and *Hspe1* have shown that chaperones play a significant role in preventing the decline in muscle mass and strength caused by disuse and certain pathological conditions. This is likely due to the fact that various chaperones are able to maintain the structure and stability of key sarcomeric proteins, preventing their aggregation, and can also influence the regulation of proteolysis and myogenesis (see review (Pomella *et al*., 2023)). More than 99% of mitochondrial proteins are encoded by the nuclear genome and synthesized in the cytoplasm; therefore, cytoplasmic and mitochondrial chaperones play a key role in the transport and import of these proteins into mitochondria. This is supported by the fact that muscle denervation-induced reduction in mitochondrial protein import occurs along with a decrease in mitochondrial HSP70 (*Hspa9*) expression and correlates with impaired mitochondrial function (Singh & Hood, 2011). This is consistent with our data on the decrease in *m. soleus* the expression of the chaperone HSPA9, the component of the inner membrane translocase TIMM50, and mitochondrial peptidases PMPCA/B, which cleave imported precursor proteins (Supporting Information Table S1). Together, these data suggest that, in our study, the larger reduction in chaperone concentrations in *m. soleus* may partially explain the more pronounced reduction in calf muscle mass and aerobic performance.

### Mitochondrial proteins

Surprisingly, no reduction in the concentration of oxidative phosphorylation enzymes, which constitute about a third of human mitochondrial proteins (Morgenstern *et al*., 2021) and are markers of mitochondrial density, was detected in *m. soleus* (Fig. A6). The literature presents mixed data on changes in mitochondrial density after 2-3 weeks of disuse (see review (Delfinis *et al*., 2025)). However, in *m. soleus*, a decrease in mitochondrial density was observed after long-term disuse (30-64 days), greater than (Hikida *et al*., 1989; Trappe *et al*., 2024) or comparable to that (Hendrickse *et al*., 2022) in *m. vastus lateralis*. Together, these findings suggest that in human skeletal muscle, disuse-induced declines in mitochondrial function occur in two stages. During the first few weeks, there is mitochondrial dysregulation, followed by a reduction in the content of respiratory enzymes and mitochondria during the second stage; moreover, *m. soleus* shows a more rapid rate of these changes compared to other muscles. Our proteomic data suggest that the dysregulation occurring in the first stage are caused by complex changes in the expression of proteins regulating proteostasis (as discussed above), mitochondrial biogenesis, oxidative stress, and calcium homeostasis (see below).

Indeed, only in *m. soleus* did disuse cause a decrease in the expression of several key regulators of mitochondrial biogenesis that control mitochondrial fusion and the formation of the mitochondrial reticulum (encoded by *MFN2* and *OPA1*), cristae morphology, the assembly and functioning of respiratory complexes (*OPA1, COA3, TMEM11*, and *LONP1*), and degradation of misfolded, unassembled, or oxidatively damaged polypeptides in the matrix (*LONP1*) (Supporting Information Table S1). The decrease in the expression of mitochondrial biogenesis regulators in “slow” *m. soleus* may explaine the greater sensitivity of this muscle to disuse compared to “mixed” *m. vastus lateralis*.

### Oxidative stress regulators

Disuse may induce oxidative stress in human skeletal muscle, causing an increase in the expression of antioxidant defense proteins (see review (Delfinis *et al*., 2025)). Our study revealed an increase in the expression of antioxidant defense proteins only in *m. soleus*: superoxide dismutase 1 – one of the key antioxidant defense enzymes, and other enzymes encoded by *TXN, GSTM2*/*P1, PARK7, PRXL2A*, etc. (Supporting Information Table S1). In addition, we observed a decrease in the expression of proteins related to the complex ‘hexokinase 2–mitochondrial non-selective channel porin 1’ (HK2-VDAC1), which is involved in the regulation of mitochondrial reactive oxygen species production (Vyssokikh *et al*., 2020) and the permeability of the mitochondrial outer membrane to ions and small molecules (Chiara *et al*., 2008). Interestingly, HK2 showed the largest magnitude of expression reduction in *m. soleus* (1.56-fold) and one of the largest in *m. vastus lateralis* (1.33-fold), whereas a decrease in VDAC1 expression was detected only in *m. soleus* (1.06-fold) (Fig. 2, Supporting Information Table S1). These results indirectly indicate oxidative stress in *m. soleus*, which may be one of the reasons for the impaired mitochondrial function in this muscle and the decrease in aerobic performance of the ankle plantar flexors.

### Calcium homeostasis regulators

Similar to the changes described above, increased expression of calcium homeostasis regulators was found primarily in *m. soleus* (Fig. 2C, Supporting Information Table S1). Disuse disrupts calcium homeostasis in muscle fibers, increasing the level of calcium ions in the cytoplasm and mitochondria (Michelucci *et al*., 2021). Therefore, in *m. soleus*, the disuse-induced increase in expression of key regulators of calcium translocation to and binding into the sarcoplasmic reticulum (encoded by *ATP2A2* [slow-twitch fiber isoform *SERCA2*] and *CASQ1*) and the decrease in expression of calcium transporters to mitochondria (encoded by *VDAC* and *MCU*) may be aimed at normalizing the elevated levels of cytoplasmic and mitochondrial calcium, respectively. Unexpectedly, disuse also increased the expression of key regulators of calcium ion release from the sarcoplasmic reticulum: Ryanodine Receptor 1 and Triadin (encoded by *RYR1* and *TRDN*), which is consistent with the increase in RYR1 expression in human *m. soleus* after ∼180-day spaceflight (Blottner *et al*., 2023). The increase in the level of these proteins may partially explain the disuse-induced (and oxidative stress-related) increase in calcium ion leakage into the cytoplasm, leading to the activation of calcium-dependent proteolysis and mitochondrial dysfunction (Michelucci *et al*., 2021). Our data indicate a disruption in the regulation of calcium homeostasis occurring predominantly in *m. soleus*, which may partially explain the greater decrease in calf muscle mass and ankle plantar flexor performance compared to the thigh muscles.

### Patterns of protein regulation depend on their function

The most significant change in the transcriptome of *m. soleus* that occur from the first days of disuse is the suppression of genes encoding regulators of energy metabolism, in particular mitochondrial oxidative enzymes (Fig. 3, Fig. A5 and (Popov *et al*., 2023; Borzykh *et al*., 2025b)). However, even after 3 weeks of disuse, most large-scale changes in the transcriptome did not translate into changes at the protein levels, for example, oxidative phosphorylation and the Krebs cycle enzymes, as well as glycolytic enzymes, regulators of lipid metabolism and muscle contraction (Fig. A6). This may be explained by the long half-life of these proteins (Fornasiero *et al*., 2018; Rolfs *et al*., 2021) and/or the insufficient duration of disuse. On the other hand, it indicates post-transcription protein buffering – a mechanism that maintain stable protein concentration despite fluctuations in mRNA concentration via the protein-specific regulation of translation (translational buffering) (Heller *et al*., 2026; Rao *et al*., 2026) and/or proteolysis (post-translational buffering) (Ishikawa *et al*., 2017). In particular, this mechanism plays a role in maintaining the stoichiometry of protein complexes (Ishikawa *et al*., 2017), which may partially explain the lack of changes in the proteins of the sarcomeres and respiratory complexes in our study. Moreover, our data show that mitochondrial protein regulation is quite specific: in contrast to mitochondrial respiratory enzymes, changes in the expression of genes encoding regulators of mitochondrial biogenesis are closely associated with changes in the level of their mRNAs (Fig. 4 and see below).

Transcriptomic and proteomic studies have shown that the level of mRNA is a poor predictor of protein expression (Liu *et al*., 2016; Buccitelli & Selbach, 2020). However, when analyzing proteins that changed expression in *m. soleus*, we found that regulation at mRNA levels played a dominant role (factor weight = 0.79) in changing the expression of some proteins (regulation pattern ‘*mRNA Down - Protein Down*’ or ‘*mRNA Up - Protein Up*’), while for others, this factor played little (factor weight = 0.19 for regulation pattern ‘*mRNA NS - Protein Down*’ and ‘*mRNA NS - Protein Up*’) or no role (regulation pattern ‘*mRNA Down - Protein Up*’) (Fig. 4B). Moreover, for the latter two cases, protein stability and unknown variables (apparently, the rate of protein synthesis) played an important role in protein regulation. Importantly, our data shows that patterns of protein regulation depend on their function. Proteins from the ‘*mRNA Down - Protein Down*’ group were enriched in functional categories associated with mitochondria. This group included key regulators of mitochondrial dynamics and structure (encoded by *MFN2, OPA1*, as well as *COA3, NDUFAF4*, and *TMEM11*), and a number of chaperones that play a key role in the transport, import, and folding of mitochondrial proteins (encoded by *HSP90AA1, HSPA9, HSPE1, HSPD1*, and *CRYAB*) (Supporting Information Table S3). In addition, for other regulatory patterns that we identified, examples of coregulation of genes with similar or complementary functions were also found: ribosomal and sarcomeric proteins were enriched in the ‘*mRNA NS - Protein Down*’ group, proteins that are part of exosomes – in the ‘*mRNA NS - Protein Up*’ group, and regulators of lipid metabolism – in the ‘*mRNA Down - Protein Up*’ group, (Fig. 4C). Coregulation of the expression of genes with similar or complementary functions is well known, including human skeletal muscle during disuse (Borzykh *et al*., 2025a). Our data show that proteins with similar or complementary functions may have a specific pattern of regulation at the mRNA, as well as at other levels of regulation. This observation complements data from studies comparing protein and mRNA stability in cells (Schwanhausser *et al*., 2011), as well as changes in protein and mRNA expression in human skeletal muscle after chronic aerobic exercise training (Makhnovskii *et al*., 2020).

### Limitations and Perspectives

There was not a control group in the study. Therefore, changes in some parameters may be partially due to factors other than disuse. To reduce the influence of these factors, we *i*) conducted a preliminary test session to familiarize participants with the MVC and maximal aerobic performance tests; *ii*) collected muscle tissue samples before all exercise tests (two weeks before the bed rest) to prevent the influence of intensive muscle contraction on investigated molecular parameters; *iii*) collected a second tissue sample the day before the end of bed rest to assess the pure effects of disuse on molecular parameters.

Transcriptomic response of mouse skeletal muscle to hindlimb suspension is sex-dependent (Tsitkanou *et al*., 2023). Here, the molecular response to disuse in different muscles was examined in males only. Therefore, comparing sex-specific transcriptomic and proteomic responses of human skeletal muscles appears to be a promising task to identify sex-specific mechanisms of the negative consequences of disuse.

Data from high-throughput methods makes it possible to reconstruct a generalized picture of regulation at different levels of cellular organization (Matsuzaki *et al*., 2021). Combining data from phosphoproteomic analysis, transcription factor activation (bioinformatics analysis), transcriptomic, proteomic, and metabolomic analysis is highly promising for studying disuse-induced changes in different muscles. This approach opens up the possibility of identifying (sex-specific) approaches to prevent/mitigate the negative consequences of disuse.

A key limitation of this study is that the effect of disuse on protein synthesis and degradation rates in different muscles and in different sample fractions (sarcomeric, mitochondrial, etc.) was not examined. Investigation of compartment-specific changes in protein turnover in the cell will help explain the changes in the disuse-induced expression of specific proteins detected in our study.

## Conclusions

Our study was the first to compare the effects of short-term (3 weeks) disuse on the phenotype, proteome, and transcriptome of the calf and thigh muscles. Disuse resulted in a greater decrease in lean mass, aerobic performance, and changes in the proteomic and transcriptomic profiles of the calf muscles/*m. soleus* than the thigh muscles/*m. vastus lateralis*. These results indicate a greater sensitivity of the calf muscles, which have a strong postural function (gravity-dependent muscles), to a decrease in contractile activity. A greater loss in calf muscle mass was associated with a decrease in the expression/deactivation of translation regulators in *m. soleus*, but not to the expression of the main sarcomeric proteins. At the same time, a significant decrease in aerobic performance of the ankle plantar flexors occurred without changing the expression of oxidative enzymes – a marker of mitochondrial density – in *m. soleus*. Our data suggest that this disorder is closely associated with changes in the expression of proteins that regulate the dynamics and structure of mitochondria, cellular and mitochondrial proteostasis, antioxidant defense, and calcium homeostasis.

Comparison of transcriptomic and proteomic data revealed that most large-scale changes in the transcriptome were not translated into changes at the protein levels. It may be due to the long half-lives of the detected proteins (predominantly highly abundant) or insufficient disuse time, and indicates the protein-specific regulation of translation (translational buffering) and/or proteolysis (post-translational buffering). On the other hand, it turned out that the pattern of regulation of proteins that changed expression depends on their function. Disuse-induced changes in the RNA levels played a dominant role in regulating expression of some proteins (such as mitochondrial biogenesis regulators), whereas for others (proteins of ribosome and exosome, etc.), this factor played little or no role. Identification of the mechanisms of this gene expression coregulation during disuse and the factors that determine the specific response of different skeletal muscles provides a foundation for developing targeted approaches to counteract the negative effects of disuse and pathologies on different muscles.

## Additional information Data availability

Raw transcriptomic data has been deposited to NCBI GEO: GSE310858. Proteomic data are available in Supporting Information Table S1. All data reported in this paper will be shared by the lead contact upon request.

## Authors’ contributions

Conceptualization: A.V.S., D.V.P.; Data curation: M.A.O., N.S.K., P.A.M., D.V.P.; Formal analysis: M.A.O., N.S.K., P.A.M., D.V.P.; Investigation: N.S.K., A.A.B., M.A.O., N.E.V., V.G.Z., R.Y.Z., T.F.V., E.M.L., A.A.P., G.V.G., V.E.N., A.V.S., D.V.P; Project administration: A.V.S., D.V.P; Resources: P.A.M., D.V.P.; Supervision: A.V.S., D.V.P.; Writing – original draft: M.A.O., N.S.K., D.V.P.; Writing – review & editing: M.A.O., N.S.K., D.V.P. All authors approved the final version of the manuscript; agree to be accountable for all aspects of the work in ensuring that questions related to the accuracy or integrity of any part of the work are appropriately investigated and resolved. All persons designated as authors qualify for authorship, and all those who qualify for authorship are listed.

## Acknowledgments

Mass spectrometry-based proteomic analysis was performed on the equipment of the “Human Proteome” Core Facility in Institute of Biomedical Chemistry (Moscow).

## References

Agarwal V, Gupta A, Chaudhary R & Kumar A. (2025). Elucidating the potential mechanism and therapeutic targets of chronic stress-induced muscle atrophy. Int Immunopharmacol 162, 115118.

Blottner D, Moriggi M, Trautmann G, Hastermann M, Capitanio D, Torretta E, Block K, Rittweger J, Limper U, Gelfi C & Salanova M. (2023). Space Omics and Tissue Response in Astronaut Skeletal Muscle after Short and Long Duration Missions. Int J Mol Sci 24.

Borzykh AA, Makhnovskii PA, Ponomarev, II, Vepkhvadze TF, Lednev EM, Rukavishnikov IV, Orlov OI, Tomilovskaya ES & Popov DV. (2025a). Transcription factors associated with regulation of transcriptome in human thigh and calf muscles at baseline and after six days of disuse. Biochim Biophys Acta Gene Regul Mech 1868, 195086.

Borzykh AA, Zhedyaev RY, Ponomarev, II, Vepkhvadze TF, Zgoda VG, Orlova MA, Vavilov NE, Shishkin NV, Lednev EM, Makhnovskii PA, Sharlo KA, Babkova AR, Vassilieva GY, Gimadiev RR, Shenkman BS, Rukavishnikov IV, Orlov OI, Tomilovskaya ES & Popov DV. (2025b). Multidirectional effect of low-intensity neuromuscular electrical stimulation on gene expression and phenotype in thigh and calf muscles after one week of disuse. Eur J Appl Physiol 125, 2431– 2447.

Brook MS, Stokes T, Gorissen SHM, Bass JJ, McGlory C, Cegielski J, Wilkinson DJ, Phillips BE, Smith K, Phillips SM & Atherton PJ. (2022). Declines in muscle protein synthesis account for short-term muscle disuse atrophy in humans in the absence of increased muscle protein breakdown. J Cachexia Sarcopenia Muscle 13, 2005–2016.

Buccitelli C & Selbach M. (2020). mRNAs, proteins and the emerging principles of gene expression control. NatRevGenet 21, 630–644.

Casuso RA, Huertas JR & Aragon-Vela J. (2024). The role of muscle disuse in muscular and cardiovascular fitness: A systematic review and meta-regression. Eur J Sport Sci 24, 812–823.

Chiara F, Castellaro D, Marin O, Petronilli V, Brusilow WS, Juhaszova M, Sollott SJ, Forte M, Bernardi P & Rasola A. (2008). Hexokinase II detachment from mitochondria triggers apoptosis through the permeability transition pore independent of voltage-dependent anion channels. PLoS One 3, e1852.

Chopard A, Arrighi N, Carnino A & Marini JF. (2005). Changes in dysferlin, proteins from dystrophin glycoprotein complex, costameres, and cytoskeleton in human soleus and vastus lateralis muscles after a long-term bedrest with or without exercise. FASEB J 19, 1722–1724.

Chopard A, Lecunff M, Danger R, Lamirault G, Bihouee A, Teusan R, Jasmin BJ, Marini JF & Leger JJ. (2009). Large-scale mRNA analysis of female skeletal muscles during 60 days of bed rest with and without exercise or dietary protein supplementation as countermeasures. Physiol Genomics 38, 291–302.

Crossland H, Skirrow S, Puthucheary ZA, Constantin-Teodosiu D & Greenhaff PL. (2019). The impact of immobilisation and inflammation on the regulation of muscle mass and insulin resistance: different routes to similar end-points. J Physiol 597, 1259–1270.

Deane CS, Piasecki M & Atherton PJ. (2024). Skeletal muscle immobilisation-induced atrophy: mechanistic insights from human studies. Clin Sci (Lond) 138, 741–756.

Delfinis LJ, Khajehzadehshoushtar S & Perry CGR. (2025). Perspectives on the interpretation of mitochondrial responses during skeletal muscle disuse-induced atrophy. J Physiol 603, 3679–3699.

Doering TM, Thompson JM, Budiono BP, MacKenzie-Shalders KL, Zaw T, Ashton KJ & Coffey VG. (2022). The muscle proteome reflects changes in mitochondrial function, cellular stress and proteolysis after 14 days of unilateral lower limb immobilization in active young men. PLoS One 17, e0273925.

Fitts RH, Trappe SW, Costill DL, Gallagher PM, Creer AC, Colloton PA, Peters JR, Romatowski JG, Bain JL & Riley DA. (2010). Prolonged space flight-induced alterations in the structure and function of human skeletal muscle fibres. J Physiol 588, 3567–3592.

Fornasiero EF, Mandad S, Wildhagen H, Alevra M, Rammner B, Keihani S, Opazo F, Urban I, Ischebeck T, Sakib MS, Fard MK, Kirli K, Centeno TP, Vidal RO, Rahman RU, Benito E, Fischer A, Dennerlein S, Rehling P, Feussner I, Bonn S, Simons M, Urlaub H & Rizzoli SO. (2018). Precisely measured protein lifetimes in the mouse brain reveal differences across tissues and subcellular fractions. Nat Commun 9, 4230.

Gram M, Vigelso A, Yokota T, Hansen CN, Helge JW, Hey-Mogensen M & Dela F. (2014). Two weeks of one-leg immobilization decreases skeletal muscle respiratory capacity equally in young and elderly men. Exp Gerontol 58, 269–278.

Hardy EJO, Inns TB, Hatt J, Doleman B, Bass JJ, Atherton PJ, Lund JN & Phillips BE. (2022). The time course of disuse muscle atrophy of the lower limb in health and disease. J Cachexia Sarcopenia Muscle 13, 2616–2629.

Haus JM, Carrithers JA, Carroll CC, Tesch PA & Trappe TA. (2007). Contractile and connective tissue protein content of human skeletal muscle: effects of 35 and 90 days of simulated microgravity and exercise countermeasures. Am J Physiol Regul Integr Comp Physiol 293, R1722–1727.

Heller EM, Barthel K, Raschle M, Schukken KM, Sheltzer JM & Storchova Z. (2026). Protein buffering of aneuploidy is driven by coordinated factors identified through machine learning. Mol Syst Biol.

Hendrickse PW, Wust RCI, Ganse B, Giakoumaki I, Rittweger J, Bosutti A & Degens H. (2022). Capillary rarefaction during bed rest is proportionally less than fibre atrophy and loss of oxidative capacity. J Cachexia Sarcopenia Muscle 13, 2712–2723.

Hikida RS, Gollnick PD, Dudley GA, Convertino VA & Buchanan P. (1989). Structural and metabolic characteristics of human skeletal muscle following 30 days of simulated microgravity. Aviat Space Environ Med 60, 664–670.

Ishikawa K, Makanae K, Iwasaki S, Ingolia NT & Moriya H. (2017). Post-Translational Dosage Compensation Buffers Genetic Perturbations to Stoichiometry of Protein Complexes. PLoS Genet 13, e1006554.

Kozlovskaya I, Sayenko IV, Sayenko D, Miller TF, Khusnutdinova DR & Melnik KA. (2007). Role of support afferentation in control of the tonic muscle activity. Acta Astronautica 60, 285–294.

Kurochkina NS, Orlova MA, Vigovskiy MA, Zgoda VG, Vepkhvadze TF, Vavilov NE, Makhnovskii PA, Grigorieva OA, Boroday YR, Philippov VV, Lednev EM, Efimenko AY & Popov DV. (2024). Age-related changes in human skeletal muscle transcriptome and proteome are more affected by chronic inflammation and physical inactivity than primary aging. Aging Cell 23, e14098.

Lambertz D, Perot C, Kaspranski R & Goubel F. (2001). Effects of long-term spaceflight on mechanical properties of muscles in humans. J Appl Physiol (1985) 90, 179–188.

Lawler JM, Song W & Kwak HB. (2006). Differential response of heat shock proteins to hindlimb unloading and reloading in the soleus. Muscle Nerve 33, 200–207.

LeBlanc AD, Schneider VS, Evans HJ, Pientok C, Rowe R & Spector E. (1992). Regional changes in muscle mass following 17 weeks of bed rest. J Appl Physiol (1985) 73, 2172–2178.

Leppek K, Das R & Barna M. (2018). Functional 5’ UTR mRNA structures in eukaryotic translation regulation and how to find them. NatRevMolCell Biol 19, 158–174.

Liu Y, Beyer A & Aebersold R. (2016). On the Dependency of Cellular Protein Levels on mRNA Abundance. Cell 165, 535–550.

Makhnovskii PA, Zgoda VG, Bokov RO, Shagimardanova EI, Gazizova GR, Gusev OA, Lysenko EA, Kolpakov FA, Vinogradova OL & Popov DV. (2020). Regulation of Proteins in Human Skeletal Muscle: The Role of Transcription. Sci Rep 10, 3514.

Mann CJ, Perdiguero E, Kharraz Y, Aguilar S, Pessina P, Serrano AL & Munoz-Canoves P. (2011). Aberrant repair and fibrosis development in skeletal muscle. Skelet Muscle 1, 21.

Marusic U, Narici M, Simunic B, Pisot R & Ritzmann R. (2021). Nonuniform loss of muscle strength and atrophy during bed rest: a systematic review. J Appl Physiol (1985) 131, 194–206.

Matsuzaki F, Uda S, Yamauchi Y, Matsumoto M, Soga T, Maehara K, Ohkawa Y, Nakayama KI, Kuroda S & Kubota H. (2021). An extensive and dynamic trans-omic network illustrating prominent regulatory mechanisms in response to insulin in the liver. Cell Rep 36, 109569.

McKee TJ, Perlman G, Morris M & Komarova SV. (2019). Extracellular matrix composition of connective tissues: a systematic review and meta-analysis. Sci Rep 9, 10542.

Michelucci A, Liang C, Protasi F & Dirksen RT. (2021). Altered Ca(2+) Handling and Oxidative Stress Underlie Mitochondrial Damage and Skeletal Muscle Dysfunction in Aging and Disease. Metabolites 11.

Mikines KJ, Richter EA, Dela F & Galbo H. (1991). Seven days of bed rest decrease insulin action on glucose uptake in leg and whole body. J Appl Physiol (1985) 70, 1245–1254.

Miotto PM, McGlory C, Bahniwal R, Kamal M, Phillips SM & Holloway GP. (2019). Supplementation with dietary omega-3 mitigates immobilization-induced reductions in skeletal muscle mitochondrial respiration in young women. FASEB J 33, 8232–8240.

Morgenstern M, Peikert CD, Lubbert P, Suppanz I, Klemm C, Alka O, Steiert C, Naumenko N, Schendzielorz A, Melchionda L, Muhlhauser WWD, Knapp B, Busch JD, Stiller SB, Dannenmaier S, Lindau C, Licheva M, Eickhorst C, Galbusera R, Zerbes RM, Ryan MT, Kraft C, Kozjak-Pavlovic V, Drepper F, Dennerlein S, Oeljeklaus S, Pfanner N, Wiedemann N & Warscheid B. (2021). Quantitative high-confidence human mitochondrial proteome and its dynamics in cellular context. Cell Metab 33, 2464–2483 e2418.

Ogneva IV, Kurushin VA, Altaeva EG, Ponomareva EV & Shenkman BS. (2009). Effect of short-term gravitational unloading on rat and mongolian gerbil muscles. J Muscle Res Cell Motil 30, 261–265.

Pavan P, Monti E, Bondi M, Fan C, Stecco C, Narici M, Reggiani C & Marcucci L. (2020). Alterations of Extracellular Matrix Mechanical Properties Contribute to Age-Related Functional Impairment of Human Skeletal Muscles. Int J Mol Sci 21.

Pomella S, Cassandri M, Antoniani F, Crotti S, Mediani L, Silvestri B, Medici M, Rota R, Rosa A & Carra S. (2023). Heat Shock Proteins: Important Helpers for the Development, Maintenance and Regeneration of Skeletal Muscles. Muscles 2, 187–203.

Popov DV, Makhnovskii PA, Zgoda VG, Gazizova GR, Vepkhvadze TF, Lednev EM, Motanova ES, Lysenko EA, Orlov OI & Tomilovskaya ES. (2023). Rapid changes in transcriptomic profile and mitochondrial function in human soleus muscle after 3-day dry immersion. J Appl Physiol (1985) 134, 1256–1264.

Puchkova AA, Shpakov AV, Baranov VM, Katuntsev VP, Stavrovskaya DM, Primachenko GK, Gorbachev VP, Tomilovskaya ES & Orlov OI. (2024). General results of a 21-day head-down bed rest study without use of countermeasures. Human Physiology 50, 808–816.

Rao S, Le AY, Persyn L & Cenik C. (2026). Translational buffering tunes gene expression in mice and humans. Genome Biol.

Ried-Larsen M, Aarts HM & Joyner MJ. (2017). Effects of strict prolonged bed rest on cardiorespiratory fitness: systematic review and meta-analysis. J Appl Physiol (1985) 123, 790–799.

Rolfs Z, Frey BL, Shi X, Kawai Y, Smith LM & Welham NV. (2021). An atlas of protein turnover rates in mouse tissues. Nat Commun 12, 6778.

Schwanhausser B, Busse D, Li N, Dittmar G, Schuchhardt J, Wolf J, Chen W & Selbach M. (2011).Global quantification of mammalian gene expression control. Nature 473, 337–342.

Senf SM. (2013). Skeletal muscle heat shock protein 70: diverse functions and therapeutic potential for wasting disorders. Front Physiol 4, 330.

Singh K & Hood DA. (2011). Effect of denervation-induced muscle disuse on mitochondrial protein import. Am J Physiol Cell Physiol 300, C138–145.

Thot GK, Berwanger C, Mulder E, Lee JK, Lichterfeld Y, Ganse B, Giakoumaki I, Degens H, Duran I, Schonau E, Clemen CS, Brachvogel B & Rittweger J. (2021). Effects of long-term immobilisation on endomysium of the soleus muscle in humans. Exp Physiol 106, 2038–2045.

Tomilovskaya E, Shigueva T, Sayenko D, Rukavishnikov I & Kozlovskaya I. (2019). Dry Immersion as a Ground-Based Model of Microgravity Physiological Effects. Front Physiol 10, 284.

Trappe TA, Minchev K, Perkins RK, Lavin KM, Jemiolo B, Ratchford SM, Claiborne A, Lee GA, Finch WH, Ryder JW, Ploutz-Snyder L & Trappe SW. (2024). NASA SPRINT exercise program efficacy for vastus lateralis and soleus skeletal muscle health during 70 days of simulated microgravity. J Appl Physiol (1985) 136, 1015–1039.

Tsitkanou S, Morena da Silva F, Cabrera AR, Schrems ER, Murach KA, Washington TA, Rosa-Caldwell ME & Greene NP. (2023). Biological sex divergence in transcriptomic profiles during the onset of hindlimb unloading-induced atrophy. Am J Physiol Cell Physiol 325, C1276–C1293.

Vikne H, Strom V, Pripp AH & Gjovaag T. (2020). Human skeletal muscle fiber type percentage and area after reduced muscle use: A systematic review and meta-analysis. Scand J Med Sci Sports 30, 1298–1317.

Vyssokikh MY, Holtze S, Averina OA, Lyamzaev KG, Panteleeva AA, Marey MV, Zinovkin RA, Severin FF, Skulachev MV, Fasel N, Hildebrandt TB & Skulachev VP. (2020). Mild depolarization of the inner mitochondrial membrane is a crucial component of an anti-aging program. Proc Natl Acad Sci U S A 117, 6491–6501.

Yu SH, Kyriakidou P & Cox J. (2020). Isobaric Matching between Runs and Novel PSM-Level Normalization in MaxQuant Strongly Improve Reporter Ion-Based Quantification. J Proteome Res 19, 3945–3954.

